# Defects in mating behavior are the primary cause of sterility in *C. elegans* males at elevated temperature

**DOI:** 10.1101/655027

**Authors:** Emily M. Nett, Nicholas B. Sepulveda, Lisa N. Petrella

**Author notes:** Department of Biological Sciences, University of Notre Dame, Notre Dame, IN, 46556.

## Abstract

Reproduction is a fundamental imperative of all forms of life. For all the advantages sexual reproduction confers, it has a deeply conserved flaw: it is temperature sensitive. As temperatures rise, fertility decreases. Across species male fertility is particularly sensitive to elevated temperature. Previously we have shown in the model nematode worm *C. elegans*, that all males are fertile at 20°C but almost all males have lost fertility at 27°C. Male fertility is dependent on the production functional sperm, successful mating and transfer of sperm, and successful fertilization post-mating. To determine how male fertility is impacted by elevated temperature we analyzed these aspects of male reproduction at 27°C in three wild-type strains of *C. elegans*: JU1171, LKC34, and N2. We found no effect of elevated temperature on the number of immature non-motile spermatids formed. There was a weak effect of elevated temperature on sperm activation that may negatively impact sperm function. In stark contrast, there was a strong effect of elevated temperature on male mating behavior and sperm transfer such that males very rarely successfully completed mating when exposed to 27°C. Therefore, we propose a model where elevated temperature reduces male fertility due to the negative impacts of temperature on the somatic tissues necessary for mating. Loss of successful mating at elevated temperature overrides any effects that temperature may have on the germline or sperm cells.

## Introduction

Survival of populations is dependent upon the ability of individuals to produce progeny. Despite the necessity for individuals to produce progeny, fertility has a flaw observed across species: it is temperature sensitive. Organisms from fruit flies and worms to sheep and humans display a dramatic loss of fertility when encountering temperatures only a few degrees higher than their optimal fertile temperature range (Harvey and Viney, 2007; Prasad et al., 2011; Rohmer et al., 2004; Wang et al., 2007). Although there is evidence that female fertility can be affected by increased temperature (Gouvêa et al., 2015; Petrella, 2014; Poullet et al., 2015; Tusell et al., 2011), male fertility is more commonly the principle cause for temperature sensitive infertility (Cameron and Blackshaw, 1980; Harvey and Viney, 2007; Petrella, 2014; Poullet et al., 2015; Prasad et al., 2011; Shefi et al., 2007; Yaeram et al., 2006). The steps and cellular pathways central to male fertility that are disrupted at elevated temperatures remain largely unknown. Here, we use the model nematode worm *Caenorhabditis elegans* to identify aspects of male fertility that are disrupted at elevated temperatures.

Classical mutant genetic screens in *C. elegans* have uncovered several temperature sensitive mutants that exhibit a lower threshold temperature at which organisms lose fertility (Conine et al., 2010; Coustham et al., 2006; Kawasaki et al., 1998). However, it is unclear if these mutants represent disruption in the pathways that are affected in wild-type organisms at elevated temperature or are an acquisition of germline temperature sensitivity through disruption of other pathways. Therefore, to understand the cellular pathways that are disrupted at elevated temperature leading to a loss of fertility in wild type organisms, experiments of wild-type animals are required. *Caenorhabditis* nematodes provide an ideal model to study temperature sensitive sterility because their optical clarity facilitates experiments designed to analyze germ cell development, sperm function *in vitro*, mating behaviors, sperm transfer, and fertilization can be measured in real time in live animals. In addition, there are >200 curated wild-type strains with genomic data that facilitate analysis of natural variation in the loss of fertility at elevated temperature (Cook et al., 2017). Loss of fertility in hermaphrodites as temperature increases has been analyzed in wild-type isolates of *C. elegans*, and the related *Caenorhabditis* species *C. briggsae* and *C. tropicalis* (Harvey and Viney, 2007; Petrella, 2014; Poullet et al., 2015; Prasad et al., 2011). In all three species, there are more thermotolerant isolates that are fertile at a temperature around 1°C above the temperature where more thermosensitive strains become sterile (Harvey and Viney, 2007; Petrella, 2014; Poullet et al., 2015; Prasad et al., 2011). For *C. elegans*, JU1171 is a more thermotolerant strain with greater than 50% of hermaphrodites fertile at 27°C; while N2 and LKC34 are more thermosensitive strains with fewer than 20% of hermaphrodites fertile at 27°C (Petrella, 2014). In *C. elegans* hermaphrodites, the primary effect of elevated temperature has been shown be the loss of functional sperm, leading to decreased self-fertility (Aprison and Ruvinsky, 2014; Gouvêa et al., 2015; Harvey and Viney, 2007; Petrella, 2014; Poullet et al., 2015; Prasad et al., 2011). There are also defects in the oogenic germline that play a role in reduced fertility at higher temperatures especially at 27°C and higher. Although it is known that both sperm and oocyte function can be affected by temperature in hermaphrodites the specific changes that occur in either gamete at elevated temperature are unknown.

Analysis of effects of elevated temperature on gametes leading to loss of fertility in hermaphrodites is complicated by the fact that they are self-fertile, producing both sperm and egg. Thus, with hermaphrodites the specific contribution of defects in either gamete to the overall loss of fertility at elevated temperature has been difficult to disentangle. Conversely, studying the effects of elevated temperature in *C. elegans* males allows for the analysis to be limited to the effects on sperm. However, male reproduction also requires that males be able to mate and successfully transfer sperm. Previously, we found that the percentage of *C. elegans* males that are fertile drops to near zero in all eight wild type strains when males were raised at 27°C (Petrella, 2014). Similar consequences to male fertility have been seen in multiple wild-type strains of *C. briggsae* and *C. tropicalis* males at elevated temperature (Poullet et al., 2015). In *C. elegans* differences between wild-type strains were seen when males were raised at elevated temperature and downshifted (27°C ➔ 20°C) in the last larval stage. Fertility was partially restored in the JU1171 thermotolerant strain after downshifting but not in LKC34 and N2 thermosensitive strains (Petrella, 2014). Unlike in hermaphrodites, the cellular or behavioral consequences of elevated temperature that lead to male loss of fertility at elevated temperature have not been investigated.

Male reproduction in *C. elegans* requires both production of haploid spermatids through the meiotic divisions of spermatogenesis and functional motile sperm through maturation during spermiogenesis. Spermatogenesis begins during larval development, in early-to-mid-L4-stage and fully formed spermatids are present by the end of the L4 larval stage (L’Hernault, 2006; Schedl, 1997). These non-motile spermatids then become activated through the process of spermiogenesis to become mature spermatozoa capable of movement and fertilization (Ellis and Stanfield, 2014). Activation of male spermatids occurs only after successful transfer to a hermaphrodite, which results in sperm motility with the formation of the pseudopod necessary for sperm crawling. Activation occurs through two redundant signaling pathways: the TRY-5 pathway in response to the TRY-5 protease in male seminal fluid and the *spe-8* pathway in response to an unknown signal in the hermaphrodite reproductive tract (Ellis and Stanfield, 2014; L’Hernault et al., 1988; Smith and Stanfield, 2011). If male spermatids do not activate properly, they cannot crawl from the uterus to the spermatheca, the hermaphrodite sperm storage organ, and will fail to fertilize an oocyte resulting in no progeny.

For males to produce progeny, sperm must be transferred to hermaphrodites through a complex male mating behavior process. These stereotypical mating behaviors involve male specific neuronal circuits and require properly formed male tail structures. The initial steps of mating require males to successfully find a hermaphrodite and subsequently locate the hermaphrodite vulva. Sensory neurons in the head allow males to find hermaphrodites in response to mating hormones (Narayan et al., 2016; Srinivasan et al., 2008; Wan et al., 2019; White et al., 2007). This is followed by signals through the nine pairs of sensory rays found in male specific tail structures, which are necessary for maintaining contact with the hermaphrodite and finding the vulva (Garcia et al., 2001; Koo et al., 2011; Liu and Sternberg, 1995; Liu et al., 2011). The final step before sperm transfer is insertion of the spicules into the vulva, which requires coordination between sensory neurons and male tail muscles (LeBoeuf et al., 2014; LeBoeuf and Garcia, 2017; Schindelman et al., 2006). Only once these mating steps are finished can males then successfully transfer sperm into the uterus. Whether loss of male fertility is due primarily to the effects of elevated temperature on the gametes, as is seen in hermaphrodites, or on somatic tissues required for mating behavior is unknown. Elevated temperature could affect one or all of these steps in males from the earliest steps of spermatogenesis to final step of sperm transfer after spicule insertion.

To determine why male *C. elegans* lose fertility at elevated temperature we analyzed male tail morphology, spermatogenesis, spermiogenesis, mating behaviors, and sperm transfer for defects at elevated temperature. Analyses were done in the more thermotolerant strain JU1171 and two thermosensitive strains LKC34 and N2. We found that there were no defects in male tail morphology or decreased sperm number in any of the three strains at 27°C males. *In vitro* sperm activation assays showed that there were no defects in activation with compounds that mimic the TRY-5 activation pathway, but there were moderate defects in activation with compounds that change ion concentrations in sperm. On the other hand, we saw large effects of temperature on both mating behavior and sperm transfer in males that experienced 27°C. Our data indicate that decreased male fertility at elevated temperature is primarily due to changes in somatic tissues that affect the ability to perform and complete mating, and that these somatic effects preclude identification of any effects that temperature may have on male sperm.

## MATERIALS AND METHODS

### Strains Used

*C. elegans* were maintained using standard procedures at 20°C on AMA1004 *E. coli* spotted NGM plates. Male strains were maintained by continually crossing males and hermaphrodites of the same genotype at 20°C. All three wild type strains used: JU1171, from Concepción, Chile, LKC34, from Madagascar, and N2, from Bristol, England and CB138 *unc-24(e138)* were obtained from the *Caenorhabditis* Genetics Center (CGC), which is funded by NIH Office of Research Infrastructure Programs (P40 OD010440).

### Temperature treatments

Four temperature treatments were used in this study. 20°C experiments were done with strains that were maintained continuously at 20°C. 27°C experiments were done by shifting P0 males and hermaphrodites to 27°C at the L4 stage and experiments were done on F1 males that had experienced their entire life span at 27°C. For both continuous 20°C and 27°C exposures, L4 males were isolated away from hermaphrodites and maintained at the experimental temperature for 24 hours prior to each analysis. For 20°C to 27°C up-shift experiments males developed at 20°C until the L4 larval stage and then were up-shifted to 27°C (in the absence of hermaphrodites) and experiments were conducted after 18 or 24 hours at 27°C. For 27°C to 20°C down-shift experiments F1 males that developed at 27°C were down-shifted to 20°C at the L4 stage (in the absence of hermaphrodites) and experiments were conducted after 24 hours at 20°C.

### Spermatid Counting

Young adult males were selected the night before experimentation and placed on AMA1004 plates without hermaphrodites. On the day of experimentation, males were transferred to NGM plates without bacteria for 5 minutes before being placed in M9 on gelatin-coated poly-L-lysine (GCP) slides (ddH_2_O, gelatin, chromium potassium sulfate, and poly-L-lysine hydrobromide) and covered with a 18mm x 18mm coverslip. Care was taken to keep from crushing males while wicking, which results in spermatids exploding from the tail. Slides were placed in liquid nitrogen for approximately 1 minute before the cover slip was removed and the slide then fixed in 100% cold methanol and acetone for 10 minutes each. Slides were incubated with block (1.5% BSA, 1.5% ovalbumin, 0.05% NaN_3_ in 1xPBS) for 30 minutes and DAPI (0.2%) for 10 minutes. Slides were washed with 1xPBS for 4×10 minutes before being mounted with a 22mm × 22mm coverslip using gelutol. The posterior ends of individual intact males were imaged in a z-stack. Images were acquired using a Leica Application Suite Advanced Fluorescence 3.2 software using Leica CTR6000 deconvolution inverted microscope with a Hamamatsu Orca-R2 camera and Plan Apo 63X/1.4 numerical aperture oil objective. Images were deconvolved. For sperm counting images of each male were imported into FIJI (Rueden et al., 2017; Schindelin et al., 2012; Schneider et al., 2012), merged into a stack, and trimmed to exclude any images within the z-stack that did not contain sperm. The hyperstacks temporal-color code function was then used to create a projection of the stack where sperm in different planes are pseudocolored different colors. The multipoint tool within FIJI was then used to count sperm until either all of the sperm within the male were counted or until at least 200 sperm were counted. Each strain was analyzed using males that were raised at 20°C or 27°C continuously. Analysis of strains was performed in two separate biological replicates for each strain from a total of ten to twelve males at each temperature condition.

### Spermatid Activation

Young adult males were selected the night before experimentation and placed on plates with a small spot of food without hermaphrodites. On the day of experimentation, males were transferred to NGM plates without bacteria prior to dissection. Spermatids were dissected from males in Sperm Buffer (5 mM HEPES pH 7.0 or 7.8, 50 mM NaCl, 25 mM KCl, 5 mM CaCl_2_, 1 mM MgSO_4_, and 10 mM dextrose) plus activator. Activators include pronase E, 200 µg/mL; triethanolamine (TEA), 60 mM; monensin, 5×10^-7^ M; 4,4’-diisothiocyano-2,2’-stilbenedisulfonic acid (DIDS), 0.4 mM; and ZnCl_2_, 1 mM. Spermatids were allowed to incubate in sperm buffer only, Pronase E, and TEA for 10 minutes, ZnCl_2_ and DIDS for 20 minutes, and monensin for 42 minutes. Spermatids were imaged on a Nikon Eclipse TE2000-S inverted microscope with Normarsky optics using Q Capture Pro 7 software with a Q imaging Exi Blue camera and Plan Apo 60X/1.25 numerical aperture oil objective. Spermatids were scored using ImageJ software as not activated (round, no pseudopod) or activated (extending a process). Three biological replicates were carried out for each activator and each strain at both 20°C and 27°C with total sperm counted between 191 and 1189.

### Male Mating Interests

Mating interest assays were performed using a protocol modified from Chatterjee et. al. (2013). Young adult males were selected the night before experimentation and placed on AMA1004 plates without hermaphrodites. On the day of experimentation, 20 CB138 *unc-24(e138)* hermaphrodites were placed on a plate with only a small area of AMA1004 bacteria and allowed to acclimate for 10 minutes. A single male was placed on this plate, and its behavior was recorded for 4 minutes (Chatterjee et al., 2013). We recorded 6 behaviors within this period. 1) Response: a positive response was recorded once the male began to scan the hermaphrodite with its tail. 2) Response time: the time at which the male began scanning the hermaphrodite. 3) Failure to Turn: a positive failing to turn response was recorded if the male reached the head or tail of the hermaphrodite but was unable to begin scanning on the opposite surface of the hermaphrodite. This appeared as the male trying to turn but eventually wandering away from the hermaphrodite. 4) Vulva located: a positive vulva located response was recorded if the male successfully found the hermaphrodite vulva while scanning and attempts to insert its spicule. 5) Number of Passes of the Vulva: the number of times the male passed over the vulva while scanning the hermaphrodite body. The LOV efficiency was calculated by vulva location/number of passes. 6) Maintain Contact: a positive maintaining contact response was recorded if the male successfully remained at the vulva during spicule insertion. Three trials of 15 males each were conducted for all strains at both 20°C and 27°C.

### Male Sperm Transfer Assays

L4 males and hermaphrodites were selected and isolated from each other for 24 hours. Young adult males were then moved to a plate spotted with 15 μl *E. coli*, and 50 μl of 0.05 mM MitoTracker Red CMXRos (Fisher Scientific cat #M7512) in 1X M9 buffer was dispensed onto the food lawn on the plate. Males were then allowed to feed for four hours. After four hours, males were subsequently moved to new plates twice to remove the residual food with MitroTracker Red. Next, adult hermaphrodites were anesthetized in a 0.1% Tricaine, and 0.01% Tetramisole in 1X M9 buffer anesthetic solution to reduce their movement and allow males to mate with them more easily. N2 worms were anesthetized for 15 minutes, JU1171 for 13 minutes and LKC34 for 10 minutes. These exposure times were used for each strain to maximize a level of anesthesia that resulted in hermaphrodites that move little but have enough muscle tone to allow for successful mating. After anesthesia, hermaphrodites were transferred to new worm plates to remove residual anesthetic solution such that anesthesia would not be encounter by males. Twelve hermaphrodites were moved to new mating plates and arranged around the food spot like a clock face. Subsequently ten males were moved to the plates with hermaphrodites. Worms were allowed to mate for 45 minutes. Each plate was monitored for insemination events, and after a hermaphrodite was inseminated it was moved to a separate plate. After the mating period, the inseminated hermaphrodites were observed for an additional 45 minutes to determine if male sperm would migrate from the uterus to the spermatheca.

### Male Tail Morphology

Young adult males were selected the night before experimentation and placed on AMA1004 plates without hermaphrodites. On the day of experimentation, males were transferred to NGM plates without bacteria prior to fixation. Fresh 2% agar pads were made daily on normal slides. Males were transferred to 10mM levamisole in 1XM9 buffer on agar pads. Male tails were imaged on the Nikon Eclipse TE2000-S inverted microscope with Normarsky optics using Q Capture Pro 7 software with a Q imaging Exi Blue camera and Plan Apo 60X/1.25 numerical aperture oil objective. Tails were scored for ray morphology (missing, fused, or swollen), spicule length, and spicule and fan smoothness. Three biological replicates of approximately 10 males were carried out for all strains at both 20°C and 27°C.

### Statistical Analysis

All statistical tests including Fisher’s Exact test and students T-test were done using Prism Graph Pad 6 or 7 software (La Jolla, CA).

## Results

### Elevated temperature did not result in low sperm count

In many species, exposure to elevated temperature can result in low sperm count in males due to effects on spermatogenesis (David et al., 2005; Wang et al., 2007; Yaeram et al., 2006). We first determined if reduced sperm count was the primary reason for lower fertility at elevated temperatures in *C. elegans*. We assessed sperm number in males raised continuously at 27°C compared to males raised at continuously 20°C. We counted sperm to determine if young adult males had at least 200 sperm (normal range), 100-200 sperm (low range), or fewer than 100 sperm (extremely low range). We found that the percentage of males raised at 27°C with a normal sperm count was the same as males raised at 20°C (Fig. 1). For both JU1171 and LKC34 males, there were a few males raised at 27°C that had a low range of sperm (100-200 sperm), but no males had fewer than 100 sperm (data not shown). Thus, like has been previously seen in *C. elegans* hermaphrodites (Harvey and Viney, 2007; Poullet et al., 2015), elevated temperature does not primarily impact male sperm count.

**Figure 1:**
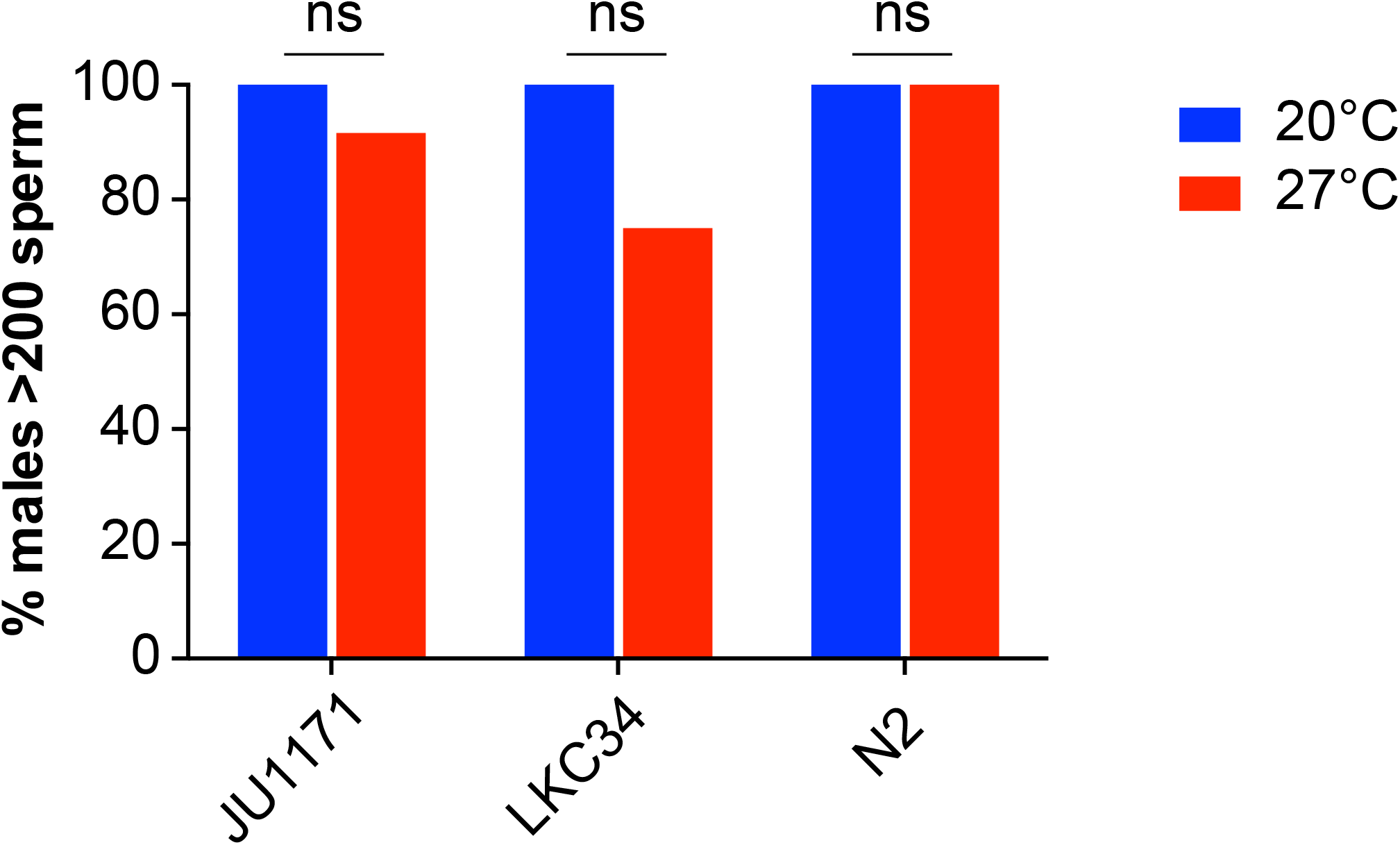
Male sperm count did not change between temperatures. Males were raised at 20°C or 27°C, and sperm were counted in intact males to determine if males had at minimum of 200 sperm as a cut-off for normal sperm number. There was no statistical difference in the number of males that had >200 sperm when raised at 20°C versus 27°C (p ≤ 0.05, Fisher’s exact test).

### *In vitro* sperm activation by zinc and DIDS is weakly affected by elevated temperature

We did not see effects of temperature on the number of spermatids indicating no significant temperature effects on the process of spermatogenesis. We next assessed effects of temperature on spermiogenesis, the formation of motile sperm through sperm maturation. Sperm are stored in males as round, non-motile spermatids (Fig. 2A). Upon ejaculation into the hermaphrodite, male sperm undergo spermiogenesis or sperm activation, forming a pseudopod resulting in motility. There are two main molecular pathways that lead to the activation of sperm; the TRY-5 pathway that is activated by the TRY-5 protease in male seminal fluid and the *spe-8* pathway that is activated by an unknown signal present in hermaphrodites (Smith and Stanfield, 2011). Using different compounds that can activate sperm *in vitro*, we assessed if these pathways were functional in sperm from males raised at 27°C continuously compared to sperm from males raised at 20°C continuously. Pronase E treatment mimics the activation of sperm through the TRY-5 pathway in the male seminal fluid (Smith and Stanfield, 2011). Pronase E treatment resulted in a small but significant decrease (9%) in activation in the JU1171 strain for sperm from males that developed at 27°C versus 20°C, but there was no difference in activation in the LKC34 and N2 strains between the two temperatures (Fig. 2B). Zinc has been suggested to activate sperm through the *spe-8* pathway that responds to signals that come from the hermaphrodite (Liu et al., 2013; Zhao et al., 2018). Zinc treatment resulted in significantly decreased activation (17-24%) in all three strains from males that developed at 27°C versus 20°C (Fig. 2B). We also tested two compounds that modify the movement of other ions in sperm, DIDS, which blocks of Clir chloride channels and monensin that causes alkalization of sperm (Machaca et al., 1996; Nelson and Ward, 1980). Which physiological activation pathway DIDS and monensin functions in is unknown. Like zinc treatment, DIDS treatment resulted in a significant decrease in activation in all three strains from males that developed at 27°C versus 20°C (Fig. 2B). Monensin treatment had mixed results between strains; in JU1171 there was significantly higher activation in sperm from males that developed at 27°C versus 20°C; in LCK34 there was no difference in activation in sperm from males that developed at 27°C versus 20°C; and in N2 there was significantly decreased activation in sperm from males that developed at 27°C versus 20°C (Fig. 2B). Overall, the data supports a model in which male growth at elevated temperature (27°C) does not affect the ability of sperm to be activated in response to the proteases in male seminal fluid as there was no or only a weak decrease in activation with Pronase E exposure. However, elevated temperature does negatively affect the ability of sperm to activate in response to changes in zinc or chloride ions.

**Figure 2:**
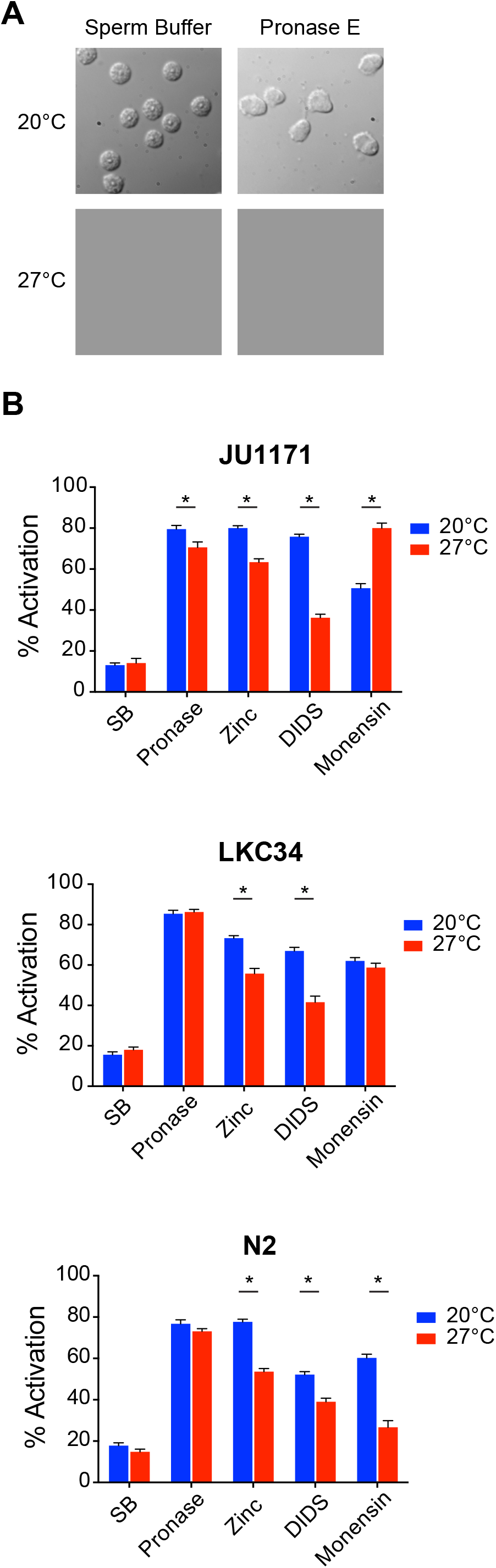
Sperm from males raised at 27°C showed decreased *in vitro* activation by induced by zinc and DIDS. (A) Sperm do not show differences in morphology from males raised at 20°C or 27°C. (B) *in vitro* sperm activation using four different activators was measured for males raised at 20°C or 27°C. Sperm from males raised at 27°C showed less activation only in JU1171 when exposed to Pronase E. Sperm from males raised at 27°C showed less activation in all three strains when exposed to zinc or DIDS. Sperm from males raised at 27°C showed less activation only in N2 when exposed to monensin (p ≤ 0.01, Fisher’s exact test) (n=191-1258). Error bars are standard error of the proportion.

### Mating behavior was substantially impacted at elevated temperature

The ability of male *C. elegans* to produce cross progeny requires both functional sperm and successful mating behavior. We wanted to assess if elevated temperature affects mating behavior thereby limiting their ability to have progeny. Stereotypical mating behavior in *C. elegans* has been well described with a number of defined steps that can be observed (Fig. 3A) (Barr and Garcia, 2006; Chatterjee et al., 2013). First, males will respond to the presence of a hermaphrodite by making contact with their tail on the cuticle of the hermaphrodite. Next, they will scan for the presence of a vulva along the length of the hermaphrodite by moving their tail along the cuticle of the hermaphrodite, often requiring the male to turn around the head or tail of the hermaphrodite. Finally, once the vulva is located the male will move back and forth near the vulva to verify the vulval location, and then maintain contact with and insert the male mating organs, the spicules, into the vulva. We observed the stereotypical mating behavior exhibited by males using a standard four-minute mating assay (Barr and Garcia, 2006) where one male is placed with a number of uncoordinated (Unc+) hermaphrodites for four minutes and monitored. We performed these assays with males exposed to two different temperature conditions during development. For continuous temperature treatments, males were raised at 20°C or 27°C throughout development. For temperature-shift treatments, males where raised at either 20°C or 27°C until the L4 stage, and then shifted to the other temperature for 24 hours. Temperature-shift experiments should bypass any effects of temperature on early development. Fewer JU1171 males completed mating when raised at 27°C continuously or when upshifted to 27°C, and specifically fewer males showed a mating response to the presence of hermaphrodites (Fig. 3B and E, Fig. S1). JU1171 males downshifted from 27°C to 20°C completed mating at levels indistinguishable from that of males raised at 20°C continuously. Fewer LKC34 males completed mating only when raised continually at 27°C. As seen in JU1171 males, LKC34 males primarily showed a decreased response to hermaphrodites when raised at 27°C continuously (Fig. 3C and E, Fig. S1). Fewer LKC34 males also showed response to hermaphrodites when downshifted from 27°C to 20°C; however, subsequent steps of mating in these males were more similar to males raised at 20°C continuously. Thus, while the overall percentage of LKC34 males that completed mating when downshifted was smaller than those LKC34 males raised at 20°C continuously, the difference was not significant. Finally, N2 males showed the strongest effects of temperature on mating behavior, with fewer males completing mating when males were exposed to 27°C during any part of their life cycle (Fig. 3E, Fig. S1). Like the other two strains, N2 also showed a decrease in the percentage of males that responded to a hermaphrodite when exposed to 27°C during development, either continuously or when downshifted. Additionally, N2 also showed a significant decrease in LOV efficiency, which is a measure of how many times the male passes the vulva before maintaining contact (Fig. 3D). A decreased LOV efficiency indicates that the male does not fully recognize the vulva and would have difficulty completing mating. Overall, males of all three strains exhibited reduced mating behavior when grown continually at 27°C and varying levels of reduced mating behavior when shifted between 20°C and 27°C. These data indicate that behavioral changes leading to a failure to mate at elevated temperature are likely the primary defect leading to male high temperature infertility.

**Figure 3:**
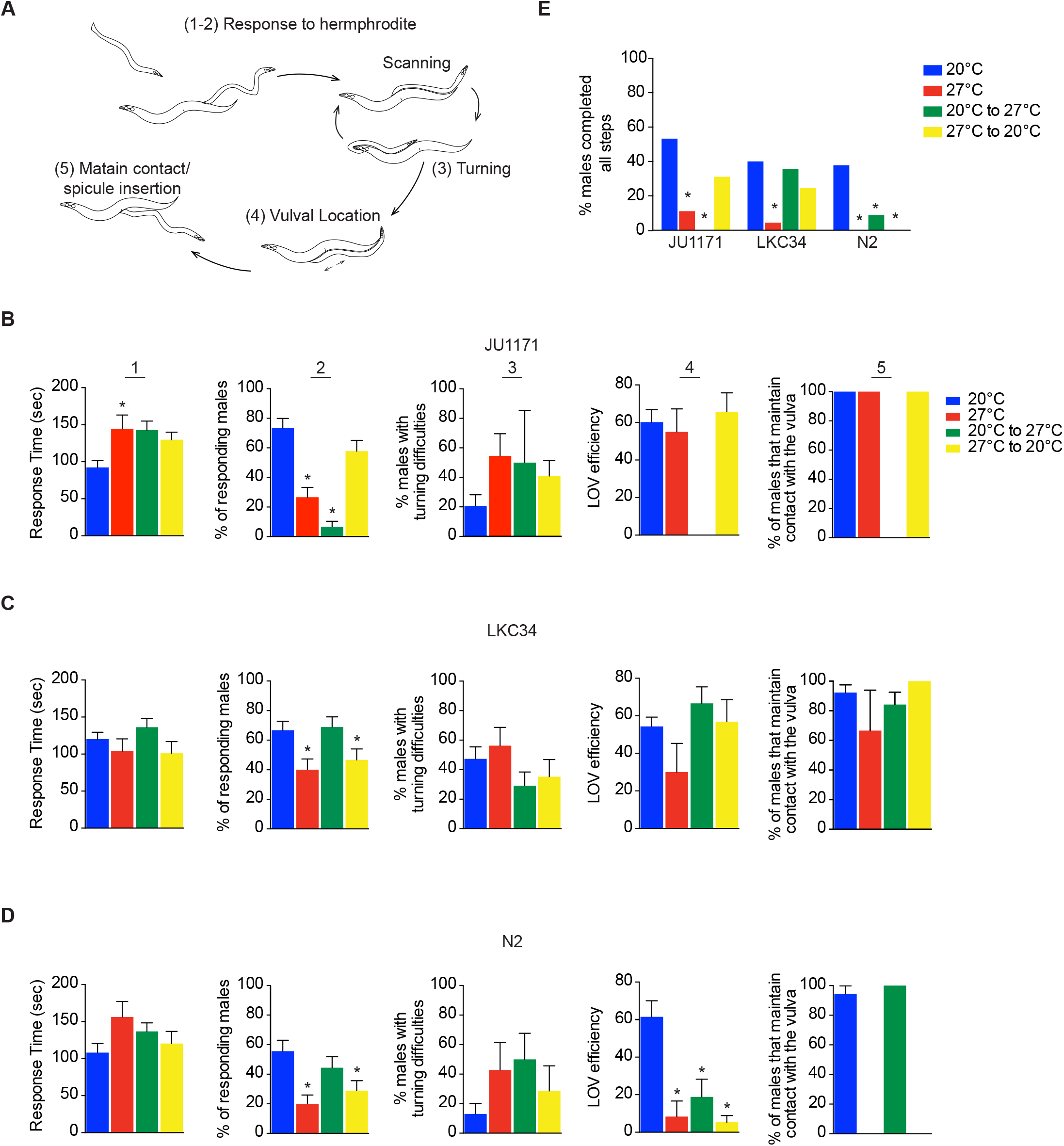
Males exposed to 27°C showed a significant decline in mating behaviors. (A) Males demonstrate stereotypical mating behaviors that include responding to the presence of hermaphrodites leading to contact (1-2), scanning along the hermaphrodite cuticle, turning if reaching the hermaphrodite head or tail (3), locating the vulva (4), and maintaining contact with the vulva (5). Redrawn from Wormbook (Barr and Garcia, 2006). (B-D) Each stage of mating behaviors was recorded for JU1171 (B), LKC34 (C), and N2 (D) strains for males raised continually at 20°C (blue), raised continuously 27°C (red), males raised 20°C then upshifted to 27°C (green), or males raised at 27°C then downshifted to 27°C (yellow). Response time (1) was increased only in JU1171 males raised at 27°C compared to JU1171 males raised at 20°C (p ≤ 0.01, Students t-test), However, there were significantly fewer males that responded to the presence of hermaphrodites (2) in all three strains, although the temperature treatment that resulted in decreased male response differed between strains (p ≤ 0.05, Fisher’s exact test). No strains showed increased turning difficulty (3) with any temperature treatment compared to males raised at 20°C (p ≤ 0.05, Fisher’s exact test). Strikingly, only the N2 strains showed a decreased LOV efficiency, which reflects increased passes across the vulva (4), but did so with any exposure to elevated temperature (p ≤ 0.01, Students t-test). Finally, if males were able to find the vulva when exposed to elevated temperature there was no significant decrease in their ability to stop movement and maintain contact (5) at the vulva (p ≤ 0.05, Fisher’s exact test). (E) As each step in the mating assay can result in the loss of a male from the mating process, we calculated the percentage of males that completed all steps successfully. LKC34 was the most temperature insensitive in this assay only showing a decreased percentage of males completing mating when males were raised continuously at 27°C. N2 was the most temperature sensitive showing decreased percentage of males completing mating with any exposure to elevated temperature (p ≤ 0.05, Fisher’s exact test). n = 15 males per data point, 3 trials. Error bars are standard error of the proportion, except for response time where error bars are standard error of the mean.

### Development at elevated temperature resulted in decreased sperm transfer

Our previous studies had indicated that males downshifted from 27°C to 20°C had very few cross progeny (Petrella, 2014), but in our mating assay both JU1171 and LKC34 downshifted males showed behaviors very similar to males kept continuously at 20°C (Fig. 3E). We next assessed if males that behaviorally complete mating also successfully transfer sperm to strain specific hermaphrodites by following the transfer of in a 45-minute window (Fig. 4A). This assay allowed us to ask a number of questions: (1) does a longer period for mating increase the number of successful matings, (2) do strain specific hermaphrodites change the number of completed matings as compared to a *unc-24* mutant hermaphrodites from an N2 background, and (3) does a male that behaviorally completed mating by maintaining contact with the vulva functionally transfer sperm to a hermaphrodite?

**Figure 4:**
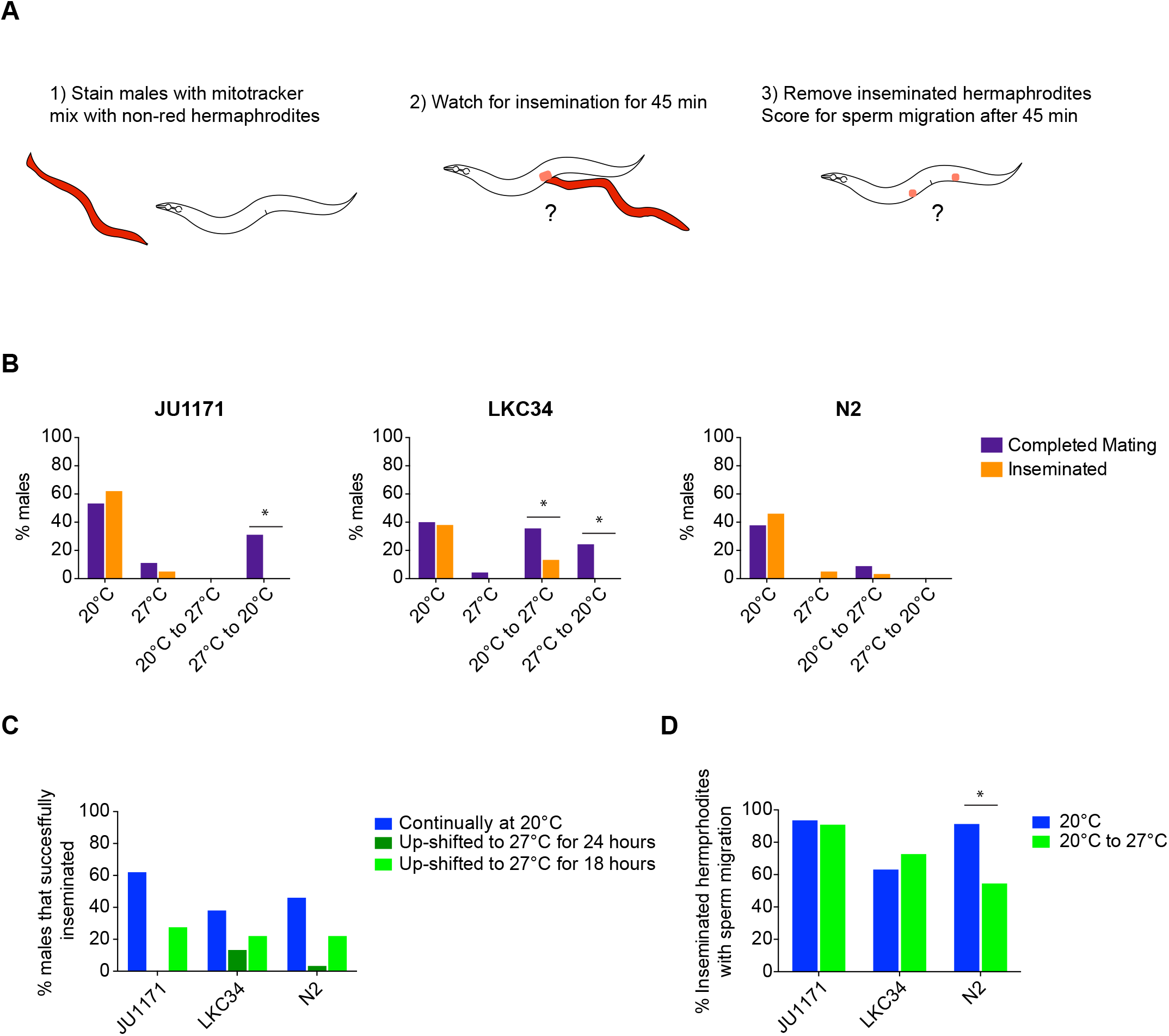
Males exposed to 27°C showed a significant decrease in sperm transfer to hermaphrodites. (A) Males were stained with MitoTracker Red CMXRos (1) and allowed to mate with anesthetized strain specific hermaphrodites for 45 min (2). Inseminated hermaphrodites were removed and scored for sperm movement to the spermatheca after 45 min (3). (B) The percentage of males that completed all steps of mating using the four min mating assay was compared to the percentage of males that could inseminate. There were significantly fewer males that could inseminate when shifted between temperatures as compared to the same temperature treatment for completing all steps of mating for JU1171 and LKC34 (p ≤ 0.05, Fisher’s exact test). (C) More males can inseminate when upshifted from 20°C to 27°C for only 18 hours instead of 24 hours. (D) The percentage of hermaphrodites that were inseminated where the sperm migrate to the spermatheca is lower for males up-shifted to 27°C for 18 hours in N2 than males kept at 20°C (p ≤ 0.05, Fisher’s exact test).

We found that when males were raised at 20°C continuously there was no significant difference between the percentage of males that could complete mating in our behavioral assay within four minutes and the percentage of males that successfully transferred sperm into a hermaphrodite within 45 minutes (Fig. 4B, Table S1). Thus, in ideal temperature conditions the behavioral assay and sperm transfer are highly correlated. Increased time did not significantly increase the rate of successful matings as assayed by sperm transfer. Additionally, all three strains were able to mate with both strain specific wild type hermaphrodites and *unc-24* mutant hermaphrodites of an N2 strain background. Under conditions in which males were raised at 27°C continuously and very few males were observed to complete mating, we also saw a very low percentage of males that could transfer sperm (Fig. 4B, Table S1). Even though both JU1171 and LKC34 males downshifted from 27°C to 20°C could behaviorally complete mating, we found that no downshifted males transferred sperm in any of the strains. Similarly, significantly fewer LKC34 males upshifted from 20°C to 27°C successfully transferred sperm than completed mating (Fig. 4B, Table S1). Overall, any exposure to 27°C resulted in significantly fewer males transferring sperm to hermaphrodites than the same strain raised at 20°C continuously. Thus, even if a male that can still behaviorally find a hermaphrodite and maintain contact with the vulva, exposure to elevated temperature limited the ability of that male to successfully transfer sperm.

### Male tail structure was not affected by elevated temperature

Successful mating and sperm transfer require male specific structures in the male tail including sensory rays and spicules. Sensory rays are needed for proper sensing of the hermaphrodite, and the spicules for penetration and ejaculation into the uterus. Since we see defects in both mating behavior and sperm transfer we analyzed male tale structure in one day old males that were raised at 20°C or 27°C continuously. We found no increase in male tails with defects in ray number or ray fusion in males raised at elevated temperature in any of the wild type isolates (Fig. S2). There was also no change in spicule length in males raised at elevated temperature (data not shown). There is no indication that there are gross defects in male tale structures at elevated temperature that should affect the ability of males to sire progeny.

### Sperm migration was affected by elevated temperature in N2

Once sperm is transferred to the hermaphrodite uterus, it must migrate to the spermatheca in order to fertilize an oocyte. Because we saw decreased *in vitro* activation levels for sperm exposed to zinc, which mimics the hermaphrodite sperm activation pathway, we wondered if male sperm exposed to elevated temperature would show decreased ability to migrate to the spermatheca. Using our fluorescent sperm transfer assay, we isolated any hermaphrodites that had received sperm and waited a minimum of 45 minutes after sperm transfer to allow for the sperm to migrate to the spermatheca (Fig. 4A). While we were able to detect migration in hermaphrodites raised at 20°C, we were limited in our ability to assess sperm migration from males exposed to elevated temperature because so few male worms mated or transferred sperm. We found that if males up-shifted from 20°C to 27°C were only exposed to 27°C for 18 hours instead of 24 hours significantly more males transferred sperm (Fig. 4C). Therefore, we compared the ability of sperm to migrate to the spermatheca from males that were raised at 20°C continuously to those males that were exposed to 27°C for ∼18 hours post-developmentally. For the two wild isolates, JU1171 and LKC34, sperm from a male exposed to 27°C for ∼18 hours migrated to the spermatheca with the same efficiency as sperm from males that had only experienced 20°C (Fig. 4D). However, for the N2 strain, sperm from a male exposed to 27°C for ∼18 hours migrated to the spermatheca significantly less often than sperm from males that have only experienced 20°C. Thus, at least for the N2 strain, even a limited exposure to 27°C can affect male sperm function within the hermaphrodite.

## Discussion

Temperature dependent defects in fertility are seen across taxa when individuals are raised outside of their optimal reproductive temperature range. Here we have completed the first analysis determining which aspects of fertility are affected by elevated temperature in *C. elegans* male. Unlike mammals and *Drosophila* (Kim et al., 2013; Rohmer et al., 2004), there is not a significant decrease in spermatid number. We also only saw moderate effects of elevated temperature on sperm activation. Instead the primary effect of elevated temperature is on mating behavior, which limits the ability of males to respond to the presence of hermaphrodites and successfully transfer sperm. These mating defects override any potential effects of temperature on sperm function. Further experiments that can circumvent the behavioral effects of temperature will be needed to determine if *C. elegans* male sperm are functional to fertilize oocytes at elevated temperature.

### Temperature effects on the soma have a large impact on fertility

One striking finding of our study is the perturbation of mating behaviors exhibited by males raised at elevated temperatures. Mating behaviors are primarily driven by interactions between somatic neurons and muscles, not by the function of the germline (Barr and Garcia, 2006). Therefore, changes to somatic tissues at elevated temperature play a large role limiting fertility in males. The primary defect we saw in males that experienced 27°C continuously was a decreased mating response compared to males exposed to 20°C continuously. We hypothesize that decreased male mating at elevated temperature is due to the temperature effects on neuronal function. The response of a male to the presence of hermaphrodites is mediated through neural input from head neurons responding to mating pheromones, including several amphid neurons and the male-specific cephalic neurons (Narayan et al., 2016; Srinivasan et al., 2008; Wan et al., 2019; White et al., 2007). The AWA neurons are one of the primary amphid neuron pairs that function in the perception of mating pheromones (Wan et al., 2019). Interesting, the AWA neurons are the sole synaptic input onto the primary neurons that sense and respond to temperature in *C. elegans*, the AFD neurons (Kimata et al., 2012; Ramot et al., 2008; White et al., 1986). Thus, temperature and mating pheromone sensing in males are part of a single circuit. Changes in temperature causes a change in the signaling rate of AFD neurons (Kimura et al., 2004; Ramot et al., 2008). This allows worms to sense and move in response to changes in temperature, a response called thermotaxis (Goodman and Sengupta, 2019; Kimata et al., 2012). The temperatures at which AFD neurons change their signaling rate leading to thermotaxis is dependent upon the temperature at which the worm developed (Clark et al., 2006; Kimura et al., 2004; Ramot et al., 2008). Therefore, the temperature to which a worm will migrate to along a temperature gradient is dependent upon the temperature at which the worm is raised (Goodman and Sengupta, 2019; Mori and Ohshima, 1995). Additionally, there is a universal increase in neuronal excitability with elevated temperature that occurs in all neurons, including both AFD and AWA neurons (Graham et al., 2008; MacIver and Roth, 1982; Ramot et al., 2008). Our data suggests that when males are raised at 27°C continuously there are changes to the neuronal circuit that senses or transmits the response to mating pheromones leading to decreased or absent mating drive. This breakdown may be due to changes in the AWA/AFD circuit where at high enough temperatures the temperature responsive nature of the AFD neurons reaches a limit and the circuit fails.

We observed defects in sperm transfer in males that were exposed to shorter treatments of 27°C. The ability to transfer sperm is primarily dependent on proper function of somatic tissues: the sensory rays in the tail must respond to the location of the vulva and the spicules must be inserted prior to sperm transfer (Barr and Garcia, 2006). Locating the vulva and insertion of spicules are dependent upon coordination between male specific tail neurons and muscles (Garcia et al., 2001; Koo et al., 2011; Liu and Sternberg, 1995; Liu et al., 2011). After spicule insertion is sensed sperm transfer will occur through a second set of coordinated neuronal signaling and muscular contraction events (LeBoeuf et al., 2014; LeBoeuf and Garcia, 2017; Schindelman et al., 2006). Increased excitability in the circuit associated with sperm transfer results in lower mating rates and an increase in early spicule insertion attempts outside the vulva (Guo et al., 2012). As temperature can also increase excitability in neural circuits (MacIver and Roth, 1982), exposure to 27°C could affect the ability of males to either insert spicules or regulate sperm transfer properly by making this circuit fire too early. In support of this possibility, we observed that males exposed to 27°C occasionally did not transfer sperm successfully into the vulva, and ejaculate was found outside the hermaphrodite on the plate (N. Sepulveda, data not shown), which we did not see with males raised at 20°C continuously. Sperm present outside the uterus can occur if sperm transfer is initiated before the completion of spicule insertion. Therefore, temperature effects on the excitability and circuit function in either the head neurons or male tail circuit could have profound effects on male mating ability.

### N2 males are more affected by elevated temperatures compared to newly isolated wild-type strains

In most of the assays we performed N2 males were more affected by elevated temperatures than either LKC34 or JU1171. N2 males showed reduced mating efficiency even with short exposure to elevated temperature, defects in locating the hermaphrodite vulva, reduced in vitro spermiogenesis upon treatment with chemical activators, and reduced sperm migration to the spermatheca. These differences may be due to the laboratory adaptation of the N2 strain in contrast to JU1171 and LKC34, which were recently isolated (Liu et al., 2013; Zhao et al., 2018) N2 is known to contain a number of unique alleles in genes that change its response to stimuli including oxygen levels and heat avoidance in comparison to recently isolated wild type strains (Andersen et al., 2014; Chang et al., 2006; Glauser et al., 2011; Sterken et al., 2015). Additionally, male mating efficiency for N2 is low compared to most other wild type isolates at 20°C, due to both a decrease in the ability of males to mate and the reception of N2 hermaphrodites to male mating (Bahrami and Zhang, 2013). The MY2 strain, which is the same isotype as JU1171, was analyzed in this study and showed approximately double the mating efficiency as N2 (Bahrami and Zhang, 2013). The long adaptation of the N2 strain in the laboratory, where self-fertilization in hermaphrodites has been selected for, may have reduced the ability of N2 males to mate even at low temperatures. N2 males may then be even more susceptible to stresses such as elevated temperature compared to males from more recently isolate strains. Careful analysis of the behavior and the effects of stressors on males from a broader range of *C. elegans* strains will be needed to determine if N2 males are a true outlier in their higher temperature sensitivity or if there is a continuum of male susceptibility to elevated temperature. However, as mating behaviors are traits that can quickly evolve (Lande, 1981; Lopes et al., 2008; Miyatake and Shimizu, 1999; Palopoli et al., 2015; Wilburn and Swanson, 2016), the lack of selective pressure to keep male mating robust in strains that are maintained in the laboratory for long periods of time seems a likely path leading to the elevated temperature sensitivity of N2 males.

### Does temperature cause a decrease in male sperm function?

In other organisms one of the major effects of elevated temperature on male fertility is a decrease in sperm count. However, in male *C. elegans* we observed that most males had a sperm count within a normal range. This mirrors what is seen in *Caenorhabditis* hermaphrodites where loss of fertility at elevated temperature is not associated with a decrease in sperm number (Harvey and Viney, 2007; Poullet et al., 2015). In *C. elegans*, although number of sperm in hermaphrodites does not decrease as temperature increases, the number of self-progeny decreases substantially (Harvey and Viney, 2007; Petrella, 2014; Poullet et al., 2015). Similarly, in *C. briggsae* and *C. tropicalis*, two related hermaphroditic *Caenorhabditis* species, there is only a small decrease in sperm number at temperatures where there is almost complete sterility (Harvey and Viney, 2007; Petrella, 2014; Poullet et al., 2015; Prasad et al., 2011). When any of these three species are raised at elevated temperature, fertility is greatly enhanced when sperm are provided from males. Therefore, the loss of fertility seen in *Caenorhabditis* hermaphrodites with elevated temperature is primarily due to a loss of sperm function, not a decrease in sperm count.

Is there a decrease in sperm function at elevated temperature in male *C. elegans* like that seen in hermaphrodites*?* While hermaphrodite and male sperm have many similarities, they also have crucial differences. Male sperm are physically bigger, there are sex-specific gene expression patterns in sperm, and there are activating factors in seminal fluid that male sperm experience which hermaphrodite sperm do not (Ebbing et al., 2018; Ellis and Stanfield, 2014; LaMunyon and Ward, 1998; Ma et al., 2014; Reinke, 2003). These differences could result in male sperm maintaining function at elevated temperature, where hermaphrodite sperm function is lost. Because of the strong effects of elevated temperature on mating behavior it is difficult to ascertain if male sperm could be functional if they were transferred into a hermaphrodite. However, there are some indications that there could be functional changes to male sperm that could contribute to decreased male fertility at elevated temperature. Using *in vitro* sperm activation assays we saw distinct differences between the ability of sperm exposed to 27°C to be activate by Pronase E and zinc or DIDS. With Pronase E there was no or only a slight decrease in *in vitro* sperm activation in sperm from males raised at 27°C. This indicates that male sperm exposed to elevated temperature are likely able to respond to and be activated by the male specific seminal fluid activation pathway through the protease TRY-5 (Fenker et al., 2014; Smith and Stanfield, 2011). However, with zinc or DIDS there was a significant decrease in activation. *In vitro* activation by zinc likely works through the *spe-8* pathway that responds to a hermaphrodite derived signal (Liu et al., 2013; Zhao et al., 2018). The recent discovery of the role of the zinc transporter protein ZIPT-7.1 highlights the important role that zinc plays in both male and hermaphrodite sperm activation(Zhao et al., 2018). ZIPT-7.1 helps maintain intracellular zinc levels within sperm and is necessary for full *in vivo* activation of sperm from both males and hermaphrodites. Classic *spe-8* pathway mutants result in hermaphrodite self-sterility, but *spe-8* pathway males are fertile unless the TRY-5 pathway is also compromised (Ellis and Stanfield, 2014). However, *zipt-7.1* mutants have highly reduced fertility in both sexes (Zhao et al., 2018). Despite the very low fertility of *zipt-7.1* mutant males, sperm from *zipt-7.1* mutant males exhibit *in vitro* activation that is only weakly reduced with Pronase E treatment compared to wild type male sperm. However, *zipt-7.1* mutant male sperm exhibit almost no *in vitro* activation with exogenous zinc treatment. The pattern we see for *in vitro* activation of sperm from males raised at 27°C looks like a dampened version of that seen in *zipt-7.1* mutants, high levels of activation with Pronase E treatment but reduced levels of activation with zinc treatment. Similar to *zipt-7.1* mutants we also see loss of both male and hermaphrodite fertility at 27°C, suggesting that changes in intracellular ion levels or fluxes, especially zinc, could lead to loss of sperm function at elevated temperatures. Additional experiments are needed to determine if male sperm that have developed at elevated temperature have defects in the *spe-8* pathway or zinc signaling.

## Acknowledgements

Some strains were provided by the CGC, which is funded by NIH Office of Research Infrastructure Programs (P40 OD010440). Many thanks to Anita Manogaran, Allison Abbott and Katherine Maniates for comments and discussion of the manuscript.

## Declaration of Interests

The authors declare no competing interests.

---

**Figure S1:**
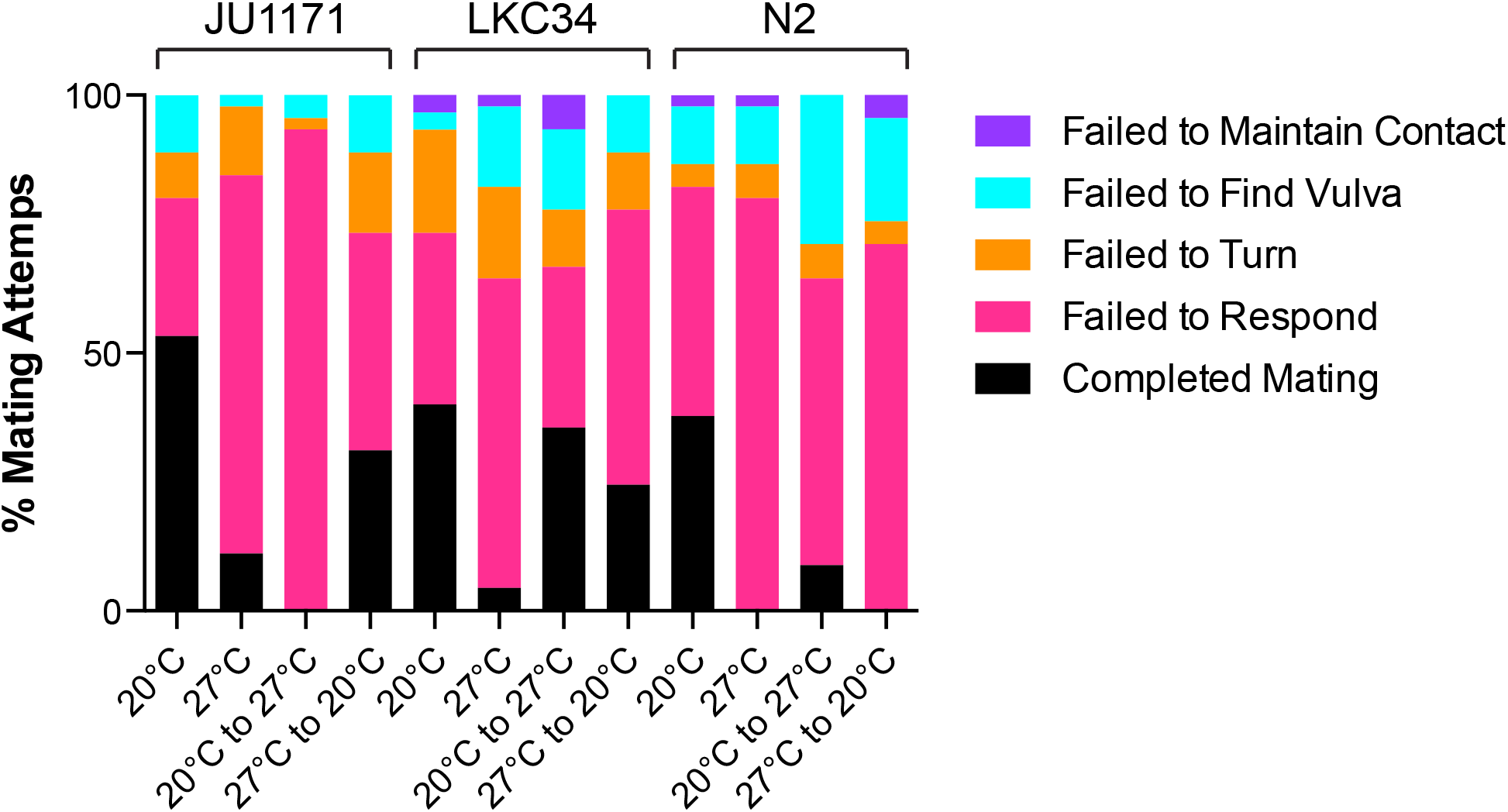
Failure to respond to the presence of hermaphrodites is most common step for males to fail in mating. Males were scored as having completed mating or designated as failed during the mating process either in not responding to the presence of hermaphrodites (Failed to Respond), not maintaining contact with the hermaphrodite after trying to execute a turn (Failed to Turn), leaving the hermaphrodite or not finding the vulva within the four minute timeframe (Failed to Find Vulva), or failing to maintain contact with the vulva once near it (Failed to Maintain Contact).

**Figure S2:**
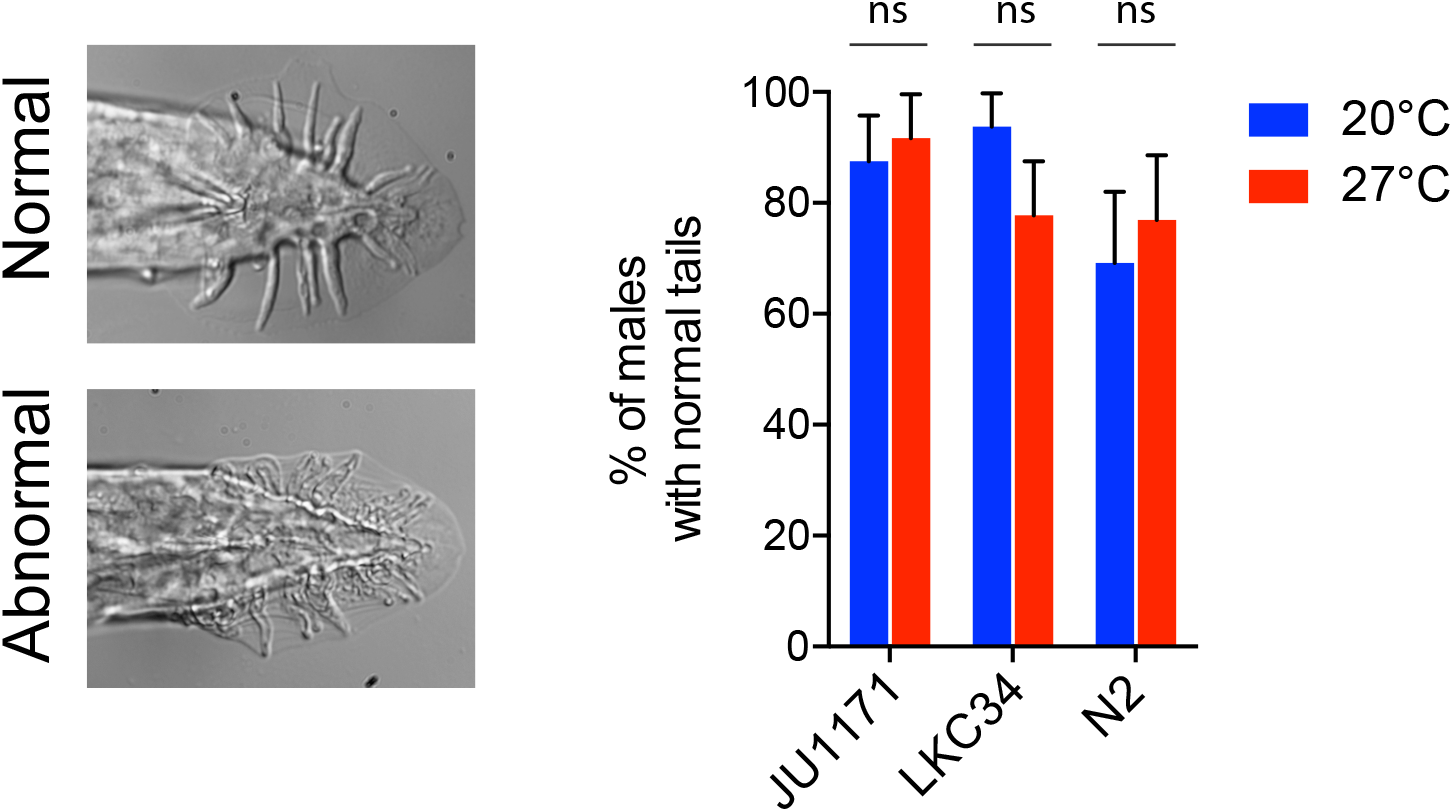
Males raised at 27°C did not show defects in tail structures. Male tails did not show increased percentage of morphological defects at 27°C. Representative images of a male tail with normal and abnormal morphology.

**Table S1:** Percentage of males that completed mating versus transferred sperm

| Strain and assay | % males in each temperature treatment |  |  |  |
| --- | --- | --- | --- | --- |
|  | 20°C | 27°C | 20°C to 27°C | 27°C to 20°C |
| JU1171 |  |  |  |  |
| Completed Mating | 53.3 | 11.1 | 0 | 31.1 |
| Inseminated | 62.0 | 5.0 | 0 | 0* |
| LKC34 |  |  |  |  |
| Completed Mating | 40.0 | 4.4 | 35.6 | 24.4 |
| Inseminated | 38.0 | 0.0 | 13.33* | 0.0* |
| N2 |  |  |  |  |
| Completed Mating | 37.8 | 0.0 | 8.9 | 0.0 |
| Inseminated | 46.0 | 5.0 | 3.33 | 0.0 |

\* p < 0.05 Fisher’s exact test between the number of male that completed mating and transferred sperm in a given strain at a given temperature

